# Genetic diversity of the eastern black rhinoceros (*Diceros bicornis michaeli*) in Tanzania; implications for future conservation

**DOI:** 10.1101/2023.01.26.525648

**Authors:** Ronald. V. K. Mellya, J. Grant C. Hopcraft, Ernest M. Eblate, Moses Otiende, Idrissa S. Chuma, Emmanuel S. Macha, Dickson Wambura, Elizabeth Kilbride, Barbara K. Mable

## Abstract

In the past decade, there has been a drastic decline in the number of Eastern Black rhinoceros (black rhinoceros) (*Diceros bicornis michaeli*), primarily because of poaching across their natural habitats, leaving few individuals in small, isolated populations that are vulnerable to demographic extinction, disease epidemics, genetic drift and inbreeding. However, genetic consequences of the demographic decline on the remaining populations have not been investigated. Using the mitochondrial control region, this study investigated how current levels of genetic diversity relate to historical patterns, quantified genetic differentiation between extant populations and assessed the impacts of previous translocations on genetic diversity across populations. A total of 74 individual eastern black rhinoceroses were sampled from five extant populations in Tanzania and one neighbouring cross-border population in the Maasai Mara in Kenya. Six maternal haplotypes were identified, with an overall haplotype diversity of h=0.7 but low overall nucleotide diversity within populations (π = 0.017) compared to historical populations from Tanzania (π = 0.021). There was extensive variation in haplotype distribution between populations, with more variation exists within (65.5 %) than among the populations (35.5%), which may indicate lack of migration between populations. Specifically, some geographically close populations with different histories of introductions didn’t share any haplotypes, suggesting that gene flow is currently restricted. The haplotypes were distributed among three east African haplogroups (CV, CE and EA) that have been described in previous studies, suggesting that multiple lineages have been preserved despite loss of haplotypes. One of the haplotypes was highly divergent and matched sequences previously classified as a subspecies that has not been recognised in recent years (*D. b. ladoensis*). We recommend that current levels of diversity be maintained by allowing natural movements of rhinoceroses between the populations, with the possibility of introducing additional variations by translocation of individuals between sites.

## Introduction

Understanding how animal populations vary within their environment is essential for developing effective conservation and managements plans; this becomes critical when dealing with endangered species. Incorporating genetic information into conservation management plans can help to reduce extinction risks by minimizing loss of genetic diversity through inbreeding, identifying populations of conservation concern, inferring population structure, resolving taxonomic uncertainties to define management units within species, detecting hybridization, defining sites for reintroductions, and choosing the best populations for reintroduction and forensics (Caughley 1994; Frankham 1995). There is, therefore, no doubt that the field of conservation genetics is key in efforts to attain sustainable biodiversity conservation.

The Eastern black rhinoceros (*Diceros bicornis michaeli*; also known as the Eastern hook-lipped rhinoceros) is a subspecies that was once widely distributed throughout South Sudan, Uganda, Ethiopia, Kenya and north-central Tanzania (Groves 1967; Kes Hillman-Smith & Groves 1994). However, the rhinoceros population has declined by 90% in the last three generations, from an estimate of 70,000 individuals across Africa in the late 1960s to only 3,800 in 1987, due to intensive poaching for their horns and habitat loss (Cumming et al. 1990). In Tanzania, the black rhinoceros population had dropped from approximately 10,000 in the 1960s to only 46 by 1997 (Brooks & Emslie 1999). The few remaining individuals were restricted to a series of small and isolated populations (Makacha et al. 1982; Sinclair & Arcese 1995). By the 1990s in Tanzania, only three populations that contained indigenous eastern black rhinoceroses remained: 1) three individuals in the Moru kopjes in the southern part of Serengeti National Park (SNP); 2) 10 individuals in the Nyamalumbwa-Maasai Mara in the northern SNP - a transboundary population between Kenya and Tanzania; and 3) 13 individuals in the Ngorongoro Crater. In 1997, conservation organizations reintroduced two female eastern black rhinoceroses to the Ngorongoro Crater from Addo Elephant National Park (Fyumagwa & Nyahongo 2010). This was followed by further reintroductions to establish new populations in Mkomazi National Park and two in the Serengeti Ecosystem (Ndasiata and Ikorongo-Grumeti) from captive populations, including: Port Lympne Wild Animal Park in the United Kingdom; Dvůr Králové Zoo in the Czech Republic; San Diego Zoo Safari Park in the United States of America; and Thaba Tholo private game ranch and Addo Elephant National Park in South Africa. Since then, these reintroduced rhinoceroses have been managed as separate subpopulations (Fyumagwa & Nyahongo 2010). As a result of the re-introductions, coupled with intensive protection and monitoring, the number of eastern black rhinoceroses in Tanzania has increased from 24 individuals in 1995 to 161 by the end of 2018 (Institute 2019). Whilst this approach has yielded success in rehabilitating these closed subpopulations, the risk of inbreeding and possible impacts of genetic drift are unknown. This is due to the fact that empirical genetic information and genetic supplementation were not considered in the selection of the founder individuals. The consequences of this demographic bottleneck on the genetic diversity in the small remote subpopulations could also result in additional impacts, including reduced viability of the population to evolve in response to extreme climates, as well as parasites and diseases (Gaines et al. 1997; 2010).

Inbreeding can put populations at risk of extinction by increasing levels of homozygosity and exposing deleterious recessive alleles that could weaken reproductive fitness and ability to survive, resulting in inbreeding depression (Frankham 1995). Furthermore, by virtue of it being stronger than selection, genetic drift can cause unpredictable loss of adaptive alleles or retention of deleterious alleles (Hartl & Clark 1997). Small, isolated populations are often also characterised by restricted gene flow, as there is less chance of immigration and emigration (Frankham 1995; Fairbairn 1998). In addition, the severity of human threats facing rhinoceroses has necessitated establishment of Intensive Protection Zones (IPZ) (Amin et al. 2006) that further restrict natural movements of individuals between subpopulations.

Apart from re-introductions from captive populations, translocation of wild individuals between different populations is another strategic management intervention (Fyumagwa & Nyahongo 2010). Such interventions are often used to balance the harmful effects of small population size and maintain natural evolutionary processes (Sinclair & Arcese 1995; Seddon et al. 2014). However, both reintroductions and translocations are only effective if the individuals being moved are sufficiently different from the host population to offset the effects of inbreeding (Jackson & Hobbs 2009). Therefore, genetic relatedness between the donor and the recipient populations is used as a key tool to inform suitability of different management interventions. It is for these reasons that establishing the current genetic health of the isolated black rhinoceros sub-populations within Tanzania and that of the neighbouring cross-border population of Maasai Mara in Kenya becomes of paramount importance.

A pan-African assessment of the genetic status of black rhinoceros populations using microsatellite markers and mitochondrial DNA (mtDNA) sequencing revealed a 69% loss of mtDNA variation of the species and identified the existence a *D. b. longipes* population in the Maasai Mara (the northern part of the Serengeti ecosystem), which had been declared extinct in 2011 (Moodley et al. 2017). They also identified sequences classified as *D. b. ladoensis*, according to the taxonomy proposed by Rookmaaker (2011); this subspecies has not always been recognised as distinct from *D. b. michaeli*. Low genetic diversity and high inbreeding were also established in the Maasai Mara sub-population on the Kenyan side compared to other larger subpopulations in a previous study (Muya et al. 2011). Furthermore, across the entire species range, seven monophyletic haplogroups have been identified (WW, west of the Shari-Logone River system; CV, east of the Shari-Logone to East Africa; NE, North-East Africa; EA, East Africa to the Zambezi River; CE, Central Africa to the Zambezi River; RU, Ruvuma region between Kilombero and Shire Rivers; SN, Southern Africa (Northern); SE, Southern Africa (Eastern); and SW, Southern Africa (Western) haplogroups (Moodley et al. 2017). Most recently, *de novo* sequence analysis of genomes from all five extant and three extinct rhinoceros species has shown strong support of the geographical hypothesis of rhinoceros evolution and confirmed low genomic diversity in all extant rhinoceroses (Liu et al. 2021). However, none of these studies included representative samples from the current populations in Tanzania, so little is known about the genetic impacts of the severe population declines and subsequent management practices to increase numbers in this region. Thus, revealing the genetic structure of the rhinoceros sub-populations will help conservation efforts with regards to current management practices focused on translocations of individuals and to inform population viability assessments.

This study used the mtDNA control region to assess the genetic diversity of eastern black rhinoceros populations in Tanzania. We aimed to address the following questions: 1) what is the current level of genetic variation in Tanzanian black rhinoceros population compared to historical estimates? 2) to what extent are the current Tanzanian subpopulations genetically differentiated from each other? and 3) how related are the extant mtDNA haplotypes in Tanzania to other haplotypes observed across Africa (i.e. how have past translocations impacted genetic diversity)?

## Materials and Methods

### Study area description

We sampled the East African subspecies of black rhinoceroses, *D. b. michaeli*, from five of the extant protected populations in Tanzania and one transboundary population in the Maasai Mara in Kenya (Figure 1). Each population has had a different history of demographic changes and re-introduction strategies as detailed below.

**Figure 1.**
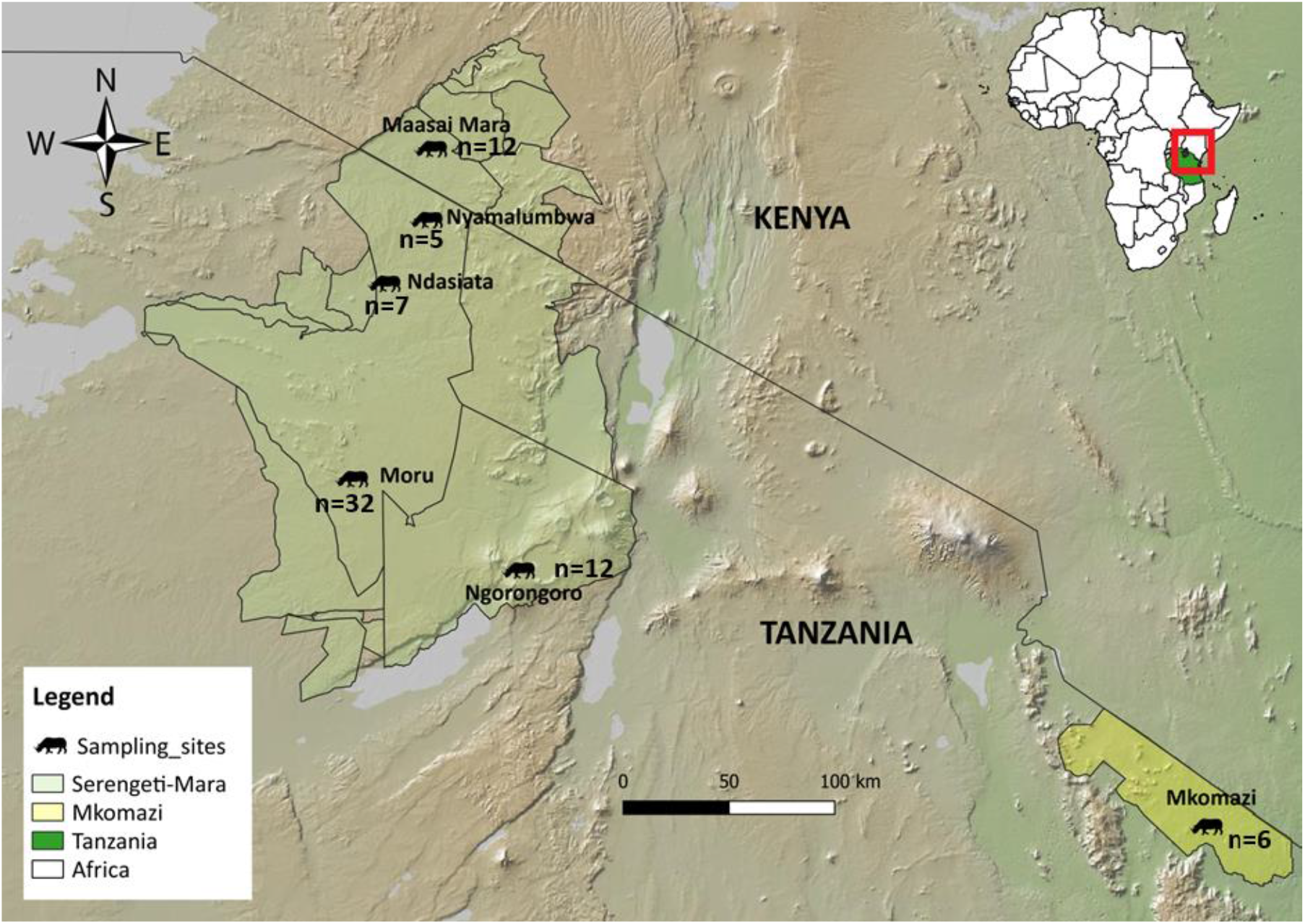
Six populations of black rhinoceroses (n=74 individuals) sampled for mtDNA analysis from the Serengeti-Mara ecosystem and the Mkomazi ecosystem in Tanzania and Kenya, East Africa. The inset shows the location of Tanzania (green) in Africa and the sampling area (in red).

#### Maasai Mara population

The Maasai Mara Game Reserve in Kenya is located in the northern portion of the Serengeti-Mara ecosystem. There were approximately 120 black rhinoceroses in 1971 but this number plummeted to 18 individuals by 1984 due to poaching (Patricia et al. 1996). It is the only population in Kenya with free-ranging indigenous inhabitants unaffected by translocations (Muya et al. 2011). At present, there are 25 rhinoceroses in this population (Table 1) that utilize areas across the border between Tanzania and Kenya and often interact with the Nyamalumbwa and Ndasiata populations within the Serengeti National Park.

**Table 1.** The number and type of samples collected from each population (n = number of individuals sequences size, N = census population size estimate based on 2018 census (TAWIRI, 2019).

| Population | Sample type |  |  |  | n | N |
| --- | --- | --- | --- | --- | --- | --- |
|  | Serum | Ear tissue | Blood EDTA | Skin biopsy |  |  |
| Maasai Mara | 0 | 12 | 0 | 0 | 12 | 25 |
| Nyamalumbwa | 0 | 5 | 0 | 0 | 5 | 20 |
| Ndasiata | 1 | 2 | 4 | 0 | 7 | 9 |
| Moru | 4 | 22 | 6 | 0 | 32 | 38 |
| Ngorongoro | 1 | 2 | 9 | 0 | 12 | 60 |
| Mkomazi | 2 | 0 | 1 | 3 | 6 | 30 |
| Total | 8 | 43 | 20 | 3 | 74 | 182 |

#### Nyamalumbwa population

The Nyamalumbwa rhinoceros project works to conserve the black rhinoceros inhabiting the cross-border area between northern Serengeti in Tanzania and the Maasai Mara National Reserve in Kenya (TAWIRI, 2019). The project started in 1999, with only four pioneer native individual rhinoceroses; i.e. one male and three females (Fyumagwa & Nyahongo 2010). The Nyamalumbwa population moves freely across the international border and often interacts with the Maasai Mara population to the North and the re-introduced population at Ndasiata to the South. There are currently 20 individuals in the population (Table 1).

#### Ndasiata population

The Ndasiata rhinoceros project (Serengeti Rhinoceros Repatriation Project) is situated in the northeastern part of the Serengeti National Park (*Figure 1*). The population was re-introduced in 2009, with the main objective to return indigenous animals to their native habitat. Five black rhinoceroses (two males and three females) were reintroduced from a captive population in Thaba Tholo, Thabazimbi, South Africa. The original animals in the Thaba Tholo captive population came from Tsavo National Park in Kenya and were caught in 1961, during a period of high poaching (Hall-Martin 1984). The population has only increased to 10 individuals (Table 1).

#### Moru kopjes population

The Moru rhinoceros project strives to conserve black rhinoceros inhabiting the southern part of the Serengeti National Park (Figure 1). The population started with only one male and two females. While the two females were residents who survived poaching crises in the late 1980’s and early 1990’s, the male migrated from the Ngorongoro Crater in 1994 (Fyumagwa & Nyahongo 2010). The three founders successfully reproduced to generate 38 individuals in the current population (TAWIRI, 2019) (Table 1).

#### Ngorongoro Crater population

The Ngorongoro Conservation Area (NCA) occupies the southern side of the Serengeti-Mara ecosystem. Between 1964-1966, there were 108 black rhinoceroses in the NCA but, due to poaching in the 1990’s, only 10 rhinoceroses remained in the area. In 1997, two female eastern black rhinoceroses were introduced from Addo Elephant National Park in South Africa. These rhinoceroses were originally taken from the Kibodo area in Kenya in 1961 and 1962 (Hall-Martin 1984). Currently, the NCA holds the largest population of free ranging rhinoceroses (Table 1) in Tanzania.

#### Mkomazi rhinoceros Sanctuary

The Mkomazi Rhinoceros Sanctuary is in Mkomazi National Park. This is actually the southern extension of Kenya’s Tsavo West National Park ecosystem (Mbeyale & Songorwa 2008) *Figure 1*. Historically, rhinoceroses would have moved between these two areas; however, fencing now restricts their movements (Homewood & Brockington 1999). The sanctuary was established in 1997 as a breeding ground for the eastern black rhinoceros population, with the aim of restoring a wild population. The starting population was composed of eastern black rhinoceros from a collection of different zoos around the world: five from Addo Elephant National Park in South Africa, three from Dvur Kravole Zoo in the Czech Republic, and three from Port Lympne Wild animal Park, UK (Fyumagwa & Nyahongo 2010). The population currently has 28 individuals (Table 1).

### Sample collection

For each population, samples were collected during ear-notching operations to provide unique individual identification and during routine veterinary interventions, and included ear tissues, whole blood in EDTA and serum. In addition, biopsy darts were used to collect tissue samples from three young individuals that had not yet been included in the ear notching campaigns (Table 1).

### DNA Extraction, mtDNA Amplification and Sequencing

Total genomic DNA from serum, EDTA blood or tissue samples was extracted using DNeasy® DNA kits following the manufacturer’s protocol (Qiagen Inc., Valencia, CA, USA, 2014). A fragment of the mtDNA control region was amplified using the mt15996L (5’-TCCACCATCAGCACCCAAAGC-3’) and mt16502H (5’-TTTGATGGCCCTGAAGTAAGAACCA-3’) primers, as described by Houlden and Brown (2000). The primers target the *D. b. michaeli* mtDNA control region at positions 15408 and 15939 (Moodley et al. 2017).

Polymerase chain reactions were carried out in a 20 μl reaction containing 2 μl of DNA diluted to 1/100, 2 μl of 1x PCR buffer, 1.2 μl of 50 mM MgCl2, 2 μl of 25 mM dNTP, 0.2 mg/μl purified BSA, 0.4 μl of each primer (10 μM), 0.2 μl of Taq polymerase (5 U/ μl) and 11.6 μl of purified water. Reactions were denatured at 95°C for 5 min, followed by 45 cycles of 94°C for 30 sec, 60°C for 1 min, 72°C for 1 min and a final extension of 72° C for 10 min. Amplified products were sent to the University of Dundee Sequencing Service for Sanger sequencing on an ABI 3730 automated sequencer; samples were sequenced in both directions using the PCR primers. The resultant sequences were manually cleaned up and the contigs assembled using Sequencher version 4.5 (Gene Codes Inc; Ann Arbor, Michigan).

### Current levels of genetic variation in Tanzania

The sequences obtained from the samples collected from extant populations were aligned with one another and grouped into unique haplotypes using Sequencher 4.5 (Gene Codes Inc; Ann Arbor, Michigan). The identity of each unique haplotype was determined using a Basic Local Alignment Search Tool (BLAST) search against the National Centre for Biotechnology Information (NCBI) database. The sequences were aligned using Clustal Omega (Sievers et al. 2011) and manually optimised using Se-Al version 2.0 (Rambaut 2002; http://tree.bio.ed.ac.uk/software/seal/). Thereafter, the sequences were collapsed into unique haplotypes using DNAsp v6 (Rozas et al. 2017) and haplotype frequencies for each population were calculated (see Supplementary Table S1).

Relationships among haplotypes were visualized using a minimum spanning haplotype network generated with PopArt version 1.7 (Leigh et al. 2015). Branch lengths were scaled according to the number of mutations separating linked haplotypes in the network.

Genetic diversity of the mtDNA control region for the entire population, as well as for each subpopulation, was independently assessed by calculating haplotype diversity (*h*) and nucleotide diversity (π) in Arlequin version 3.5 (Excoffier & Lischer 2010). Haplotype diversity (*h*) is the probability that two randomly sampled haplotypes from a population will be different from one another (Nei 1987). Nucleotide diversity (π) is the average number of nucleotide differences per site between two DNA sequences across all possible pairs in the sample population (Nei 1987). To assess changes in diversity over time, we compared the values from the extant Tanzanian and Maasai Mara populations to the historical populations sampled by Moodley et al (2017).

### Differentiation between subpopulations in Tanzania

For the current Tanzanian mtDNA control region sequences, population structure was assessed using a hierarchical analysis of molecular variance (AMOVA) in Arlequin 3.5 software (Excoffier & Lischer 2010). A matrix of genetic distances among sampled individuals was used to test specific hypotheses about the sources of variation estimated at various hierarchical levels within vs between populations to infer population differentiation based on fixation indices (i.e. F-statistics). Population differentiation was further assessed using pairwise genetic distances between each population based on F_st_, as implemented in Arlequin 3.5 software (Excoffier & Lischer 2010). The pairwise F_st_ values for each subpopulation was given in the form of a matrix to compare populations and their standardised pairwise differences between populations. We tested the significance of the F_st_ values by creating a null distribution under the hypothesis of no difference between the subpopulations by permuting (1000) haplotypes between populations and testing the proportion of permutations leading to a F_st_ value larger or equal to the observed one.

### Phylogenetic context of Tanzanian haplotypes

For comparative analysis, mtDNA D-Loop data from captive and wild eastern black rhinoceros populations were obtained from GenBank. These sequences were deposited by Moodley et al. (2017), Muya et al. (2011) and Githui et al. (2014) (see Supplementary Table S1). We used the ClustalW multiple alignment package in the BioEdit software version 7 (Alzohairy 2011) to align sequences obtained from the current study with a total of 423 other sequences retrieved from GenBank. The sequences were then collapsed into unique haplotypes and their frequencies recorded. The geographical region for each sample was identified (where that information was available; supplementary Table S1) and each haplotype classified into one of the seven monophyletic haplogroups identified by Moodley et al (2017). Where possible, haplotypes were further classified into either historical or modern groups (i.e. originating from museum archives, as opposed to being sampled from an extant population).

Phylogenetic relationships among the haplotypes was analysed by reconstructing an intraspecific phylogeny tree from black rhinoceros haplotypes recovered in our data set with a white rhinoceros (*Ceratotherium simum simum*) sequence from GenBank as an outgroup (FJ004916.1; Supplementary Table S1). this analysis was done in BEAST v 2.5 (Bouckaert et al. 2019) under a Bayesian skyline model for lineage coalescence and TN93 nucleotide substitution model, as determined by model selection in MEGA X software (Kumar et al. 2018). The analysis was run for 100 million MCMC steps sampling the posterior distribution every 10,000 steps. The initial 10% of steps were discarded to ensure we sampled from the stationary part of the distribution. The final tree was visualised in Evolview software version 3 (Subramanian et al. 2019) and annotated using: the relative frequency of each haplotype, whether the haplotype was sampled from the extant (modern) or historical (museum samples) populations, the geographical regions of the haplotypes, and the seven monophyletic haplogroups (WW, NE, CV, EA, CE, RU and SN) described by Moodley et al. (2017). For our samples and those lacking spatial data from other studies, we assigned haplogroups based on the positions in the phylogenetic tree (Supplementary Table S1).

## Results

### Current levels of genetic variation in Tanzania

A total of 62 samples were sequenced successfully from Tanzania, with 12 more sequenced from the Maasai Mara and four from Lake Nakuru in Kenya. The sequences included 25 polymorphic sites with no insertions or deletions and 438 monomorphic sites. Six haplotypes were found among the samples, which differed in frequency and distribution among the populations (Figure 2). A comparison of the sampled mtDNA haplotypes with sequences in GenBank showed close similarity to published sequences from *D. b. michaeli* from Kenya for all the haplotypes except one (Haplotype 2), which most closely matched a sequence from *D. b. lodoensis* from Uganda (Table 2). Mkomazi, Maasai Mara and Ngorongoro had the highest number of haplotypes (three each) while Moru and Ndasiata had two each and Nyamalumbwa had single haplotype. Haplotype 1 was found at the highest frequency and was shared among the four native populations from Moru, Ngorongoro, Nyamalumbwa and Maasai Mara. Haplotype 3 and 6 were shared among the two populations (Mkomazi and Ndasiata) that were formed from translocated individuals, although the latter was also found in Ngorongoro, which also contains some translocated individuals. Haplotype 2 was found only in the native Moru and Maasai Mara populations, haplotype 4 was restricted to Ngorongoro and Maasai Mara while haplotype 5 was found in the reintroduced Mkomazi population only (Figure 2). The Lake Nakuru population had two haplotypes (33 and 42) that had been found previously in that region in previous studies (Githui et al. 2014; Moodley et al. 2017; Muya et al. 2011).

**Figure 2.**
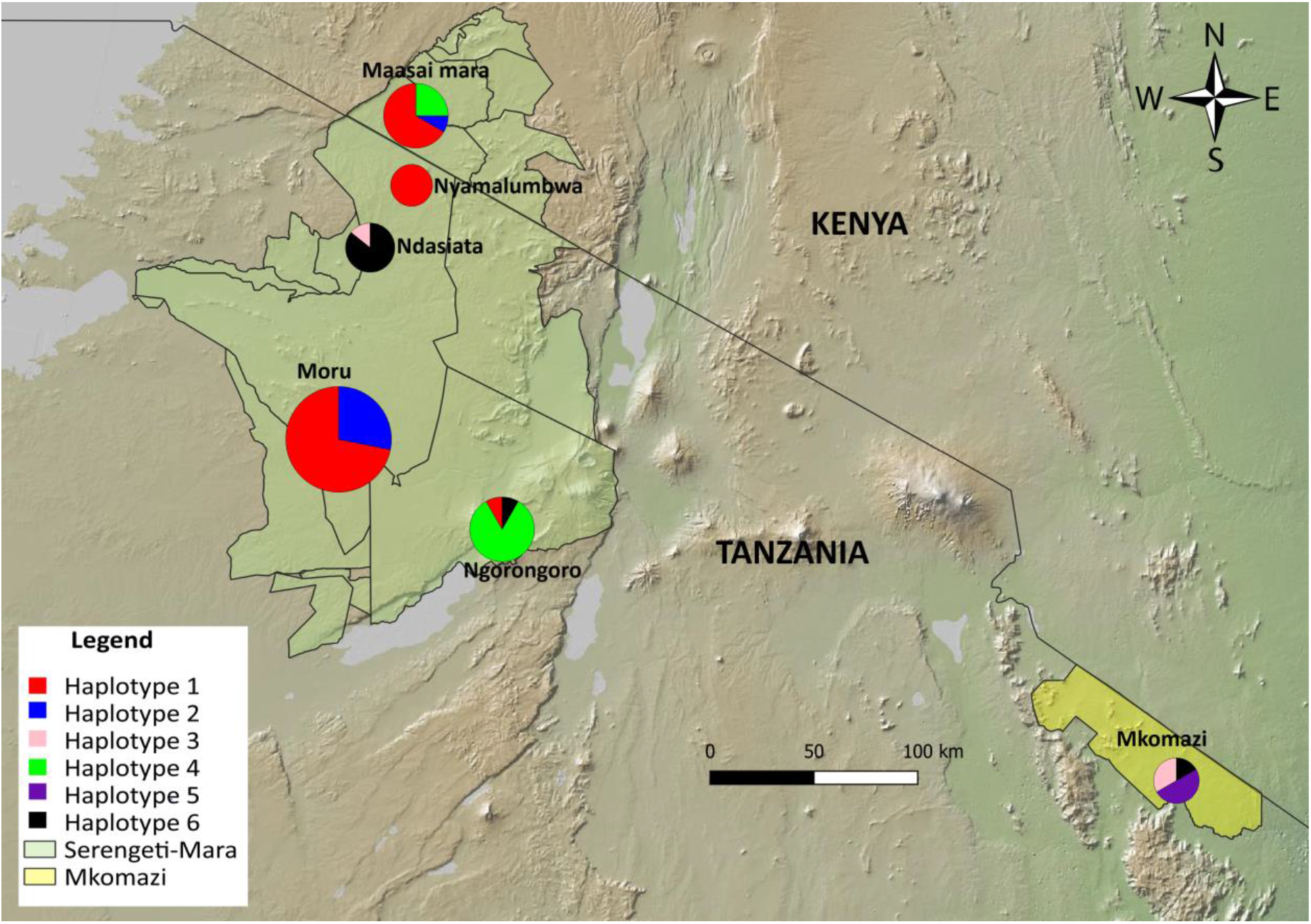
A map of relative frequency and geographical distribution of the six mtDNA haplotypes in six black rhinoceros populations in Tanzania and Kenya. Size of the circles correlates with the number of individuals sampled from each population.

**Table 2.** NCBI Blast results of the six mtDNA control region haplotypes from five extant black rhinoceros populations in Tanzania, showing the most similarly matching sequence in the GenBank database (100% similarity in each case). Query cover % = the percentage of overlap between the input sequence and the sequences identified in the database; N Match = the number of identical base pairs divided by the query length. The species identity and geographical origin of the closest match, along with the GenBank accession number are also shown.

| Haplotype | Query cover % | N Match | Species | Origin | GenBank accession |
| --- | --- | --- | --- | --- | --- |
| Haplotype 1 | 99 | 460/460 | *D. b. michaeli* | Lake Nakuru, Kenya | KP247508.1 |
| Haplotype 2 | 93 | 435/435 | *D. b. ladoensis* | Uganda | KY472411.1 |
| Haplotype 3 | 99 | 460/460 | *D. b. michaeli* | Lake Nakuru, Kenya | KP247519.1 |
| Haplotype 4 | 93 | 435/435 | *D. b. michaeli* | Ngorongoro, Tanzania | KY472537.1 |
| Haplotype 5 | 99 | 435/435 | *D. b. michaeli* | OI Pajeta, Kenya | KY472503.1 |
| Haplotype 6 | 99 | 435/435 | *D. b. michaeli* | OI Pajeta, Kenya | KY472414.1 |

The minimum spanning network showed that haplotype 2 had the highest number of mutations separating it from all other haplotypes, whereas haplotype 6 and haplotype 3 were separated by only one mutation (Figure 3). Haplotype 4 was separated from haplotype 1 and haplotype 5 by the same number of mutations; hence, the three haplotypes formed a network circle (Figure 3).

**Figure 3.**
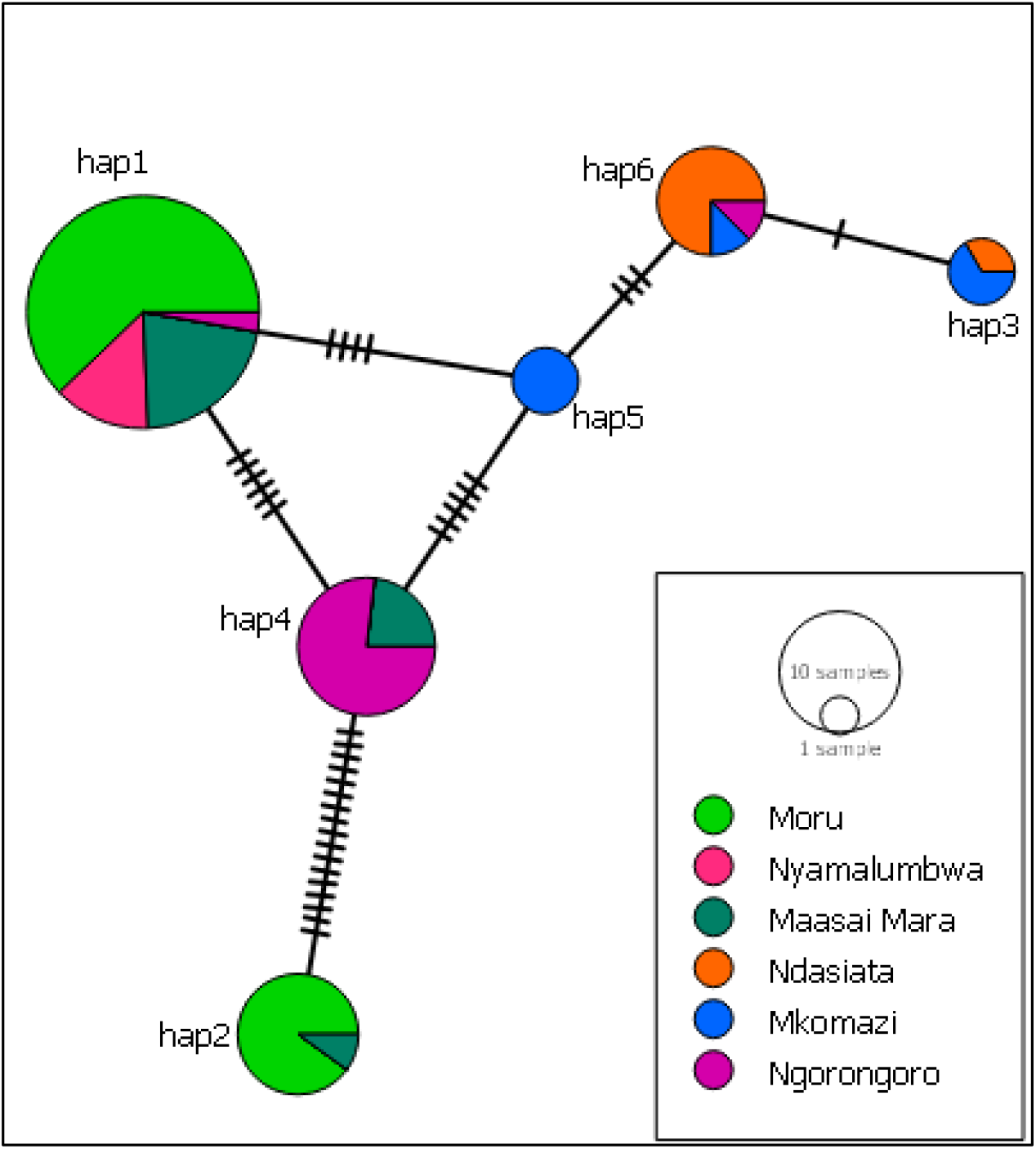
A minimum spanning network joining the six mtDNA control region haplotypes found in Tanzania. Circles represent haplotypes and the size is proportional to the haplotype frequency; ticks on branches show the number of mutations separating linked haplotypes; colours indicate the relative frequency of the haplotypes in each population.

The overall mtDNA haplotype diversity across all extant black rhinoceros sampled (n = 74) was 0.7, but the values varied considerably when each population was considered alone (Table 3). Mkomazi (n=6) had the highest haplotype diversity (0.73), while Nyamalumbwa had no haplotype diversity because it had only a single haplotype. Despite having only two haplotypes, Moru had the highest nucleotide diversity (π = 0.018), followed by Maasai Mara (π = 0.012); they shared the highly divergent haplotype 2 (Table 3).

**Table 3.** Mitochondrial DNA control region diversity of the current black rhinoceros populations in Tanzania and the Masaai Mara compared to historical samples described by Moodley et al. (2017). n=number of individuals sampled; nhap=number of haplotypes; S = number of segregating sites; h = haplotype diversity; π = nucleotide diversity. Historical diversity estimates were obtained from (Moodley et al. 2017).

| Population | n | nhap | S | h | $\pi$ |
| --- | --- | --- | --- | --- | --- |
| Maasai Mara | 12 | 3 | 21 | 0.53 | 0.012 |
| Nyamalumbwa | 5 | 1 | 0 | 0 | 0.000 |
| Ndasiata | 7 | 2 | 1 | 0.28 | 0.001 |
| Moru | 32 | 2 | 19 | 0.42 | 0.018 |
| Ngorongoro | 12 | 3 | 9 | 0.32 | 0.004 |
| Mkomazi | 6 | 3 | 4 | 0.73 | 0.005 |
| Total Extant | **74** | **6** | **25** | **0.7** | **0.017** |
| Tanzania historical | **29** | **19** | **36** | **0.95** | **0.021** |

The 19 haplotypes identified among 29 individuals sampled from historical populations in Tanzania by Moodley et al. (2017) included five of the haplotypes found in the current populations (Table 3). Haplotype 5 was not found among the historic samples from Tanzania but it had been identified among recent Kenyan samples (n=4) and a Ugandan historic sample in Moodley et al. (2017). Of the three Tanzanian samples that Moodley et al. (2017 classified as “modern”, two had haplotype 4 and one had an additional haplotype not found in our extant samples (Supplementary Table S1). Moodley et al. (2017) identified three haplotypes from eight “modern” individuals from the Maasai Mara (all collected in 1989); however, they did not find haplotype 2 in this population. Instead, they found an additional haplotype that was not found in the current samples analyzed by this study (hap63; Supplementary Table S1). Haplotype diversity for Moodley’s historical samples from Tanzania was higher (h=0.95) than for current populations in this study (h=0.73; Table 3). The average nucleotide diversity across populations in the current study was less than (π = 0.017) of the previously described historical samples from Tanzania (π = 0.021).

### Differentiation between subpopulations in Tanzania

For the comparison using AMOVA analyses, substantially more variation was explained within populations than for the higher levels of comparison but there was no evidence of significant population differentiation (Table 4). The results show that more variation exists within (65.5 %) than among the populations (35.5%), which may indicate lack of migration between populations. Furthermore, based on the pairwise F_st_ between the populations, high differentiation was observed between Ndasiata-Nyamalumbwa rhino population (Figure 4) even though being very close geographically. This might be due to isolation and limited gene flow with other populations

**Table 4.** Partitioning of genetic variation using analyses of molecular variance (AMOVA) for different levels of comparisons. The fixation index F_st_ measures the proportion of genetic variation occurring among groups. The % variation is the amount of diversity in the population associated to the partitioned group. Significance of F_st_ was assessed using 1,000 permutations.

| Comparison | Source of variation | % Variation | Variance component | p value |
| --- | --- | --- | --- | --- |
| All populations | Among populations | 35.5 | 2.59 | 0.0001 |
|  | Within populations | 64.5 | 1.43 |  |

**Figure 4.**
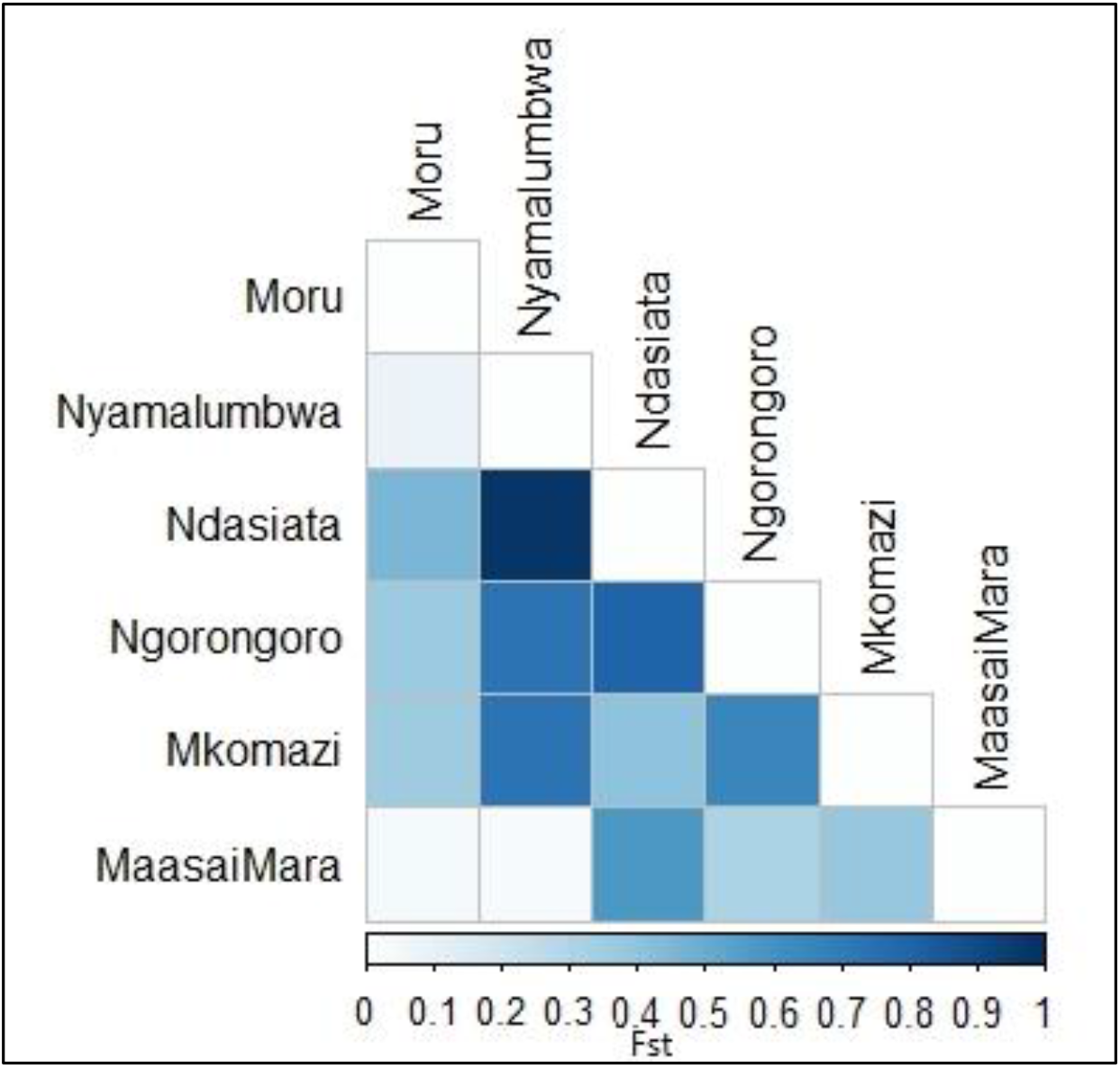
Matrix of the pairwise F_st_ (based on F-Statistics) between the Eastern black rhinoceros’ subpopulation in Tanzania. Increasing colour intensity in the heatmap reflects a higher F_st_ value. F_st_ values range from 0 for no differentiation to 1.0 for complete differentiation among subpopulations fixed for different alleles.

### Phylogenetic context of Tanzanian haplotypes

Based on a search of Genbank, no additional Tanzanian samples were available other than those described in Moodley et al. (2017); all six of the Tanzanian haplotypes identified here had been found in other East African populations in that study. However, two additional studies reported samples from Kenya (Muya et al. 2011; Githui et al. 2017) but without the date of sampling, so these were considered as “unclassified”. This allowed a more extensive assessment of the relative frequency of East African haplotypes in a broader context (Supplementary Table S1; Figure 5). Among the 144 sequences available from Kenya, 35 haplotypes were found, with 11 among samples classified as recent, 8 identified only in historic samples, and two unclassified. Haplotype 1 was found at a substantially higher frequency in Kenya than all other haplotypes (n = 30; 21% of samples); the next most frequent were haplotype 33 (n = 15; not found in Tanzania) and haplotype 3 (n = 14; found only in the translocated populations Ndasiata and Mkomazi in Tanzania). All three of these haplotypes were found in historic, recent and unclassified samples from Kenya but haplotype 3 was not found in the Maasai Mara; as in our study, Moodley et al. (2017) found haplotypes 2 and 4 in modern samples from that population, but with an additional haplotype (haplotype 63) that we did not identify. Haplotype 2, which matched D. *b. ladolensis*, was also found in historical populations from Uganda. Haplotype 6, which was found in the reintroduced populations (Ndasiata, Ngorongoro and Mkomazi) from South Africa and Europe, was also detected in Tanzanian historical populations and modern, historic and unclassified Kenyan populations. The other haplotype (haplotype 5) was not detected in historical samples from Tanzania but was found in Kenyan modern populations.

**Figure 5.**
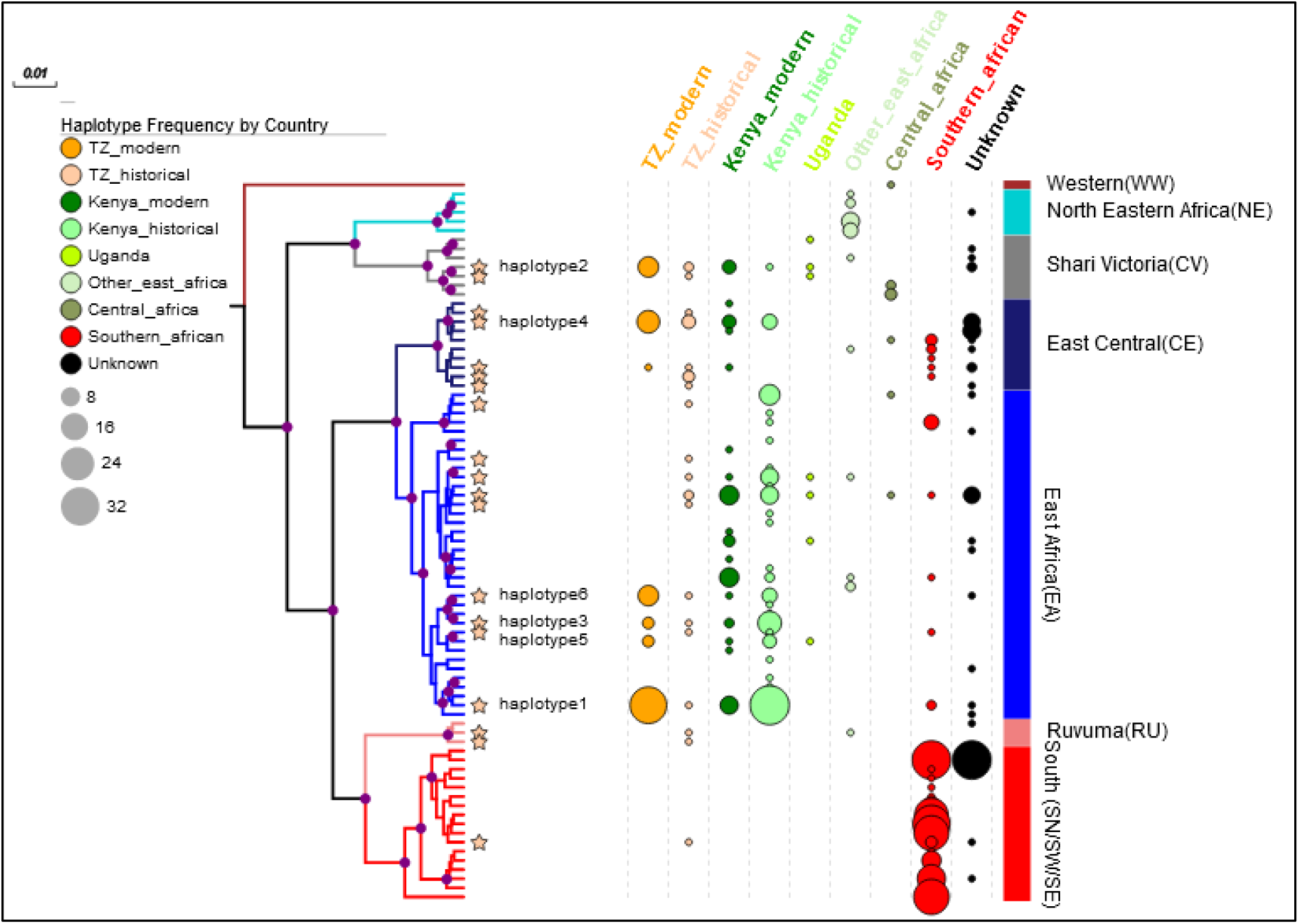
Bayesian phylogenetic tree of 79 mitochondrial DNA (mtDNA) control region haplotypes, obtained from a sample of 517 individual black rhinoceros sequences, with white rhinoceros (Ceratotherium simum simum) used as an outgroup. Branches with a posterior probability of 85% are indicate with purple node. The size of the circles correlates with the frequency of each haplotype in its geographical region (indicated by colors). Stars signify haplotypes from Tanzania, with the yellow circles indicating relative frequency in modern samples and orange circles historic. Relative frequency of haplotypes from other East African population (Kenya, Uganda and others) are indicated in shades of green. Haplotypes from individuals sampled from southern Africa are indicated in red. Note that haplotypes that had been translocated to Tanzania from South African captive populations (haplotype 3, haplotype 5, haplotype 6) all were found among East African historic samples.

Phylogenetic reconstruction of the mtDNA haplotypes using all published sequences (Figure 5) showed three divergent lineages (using the classifications described in Moodley et al. (2017)), the most distinct of which (L1) comprised haplotypes sampled from West Africa (haplogroup WW from Nigeria and Cameroon). The second lineage (L2) was separable into two haplogroups: North-eastern (NE) and North-western African (CV). The last lineage (L3) is broadly distributed in eastern and southern sub-Saharan Africa, forming six monophyletic haplogroups: EA, Eastern Africa; CE, Eastern Africa (Central); RU, Ruvuma (Eastern Africa South); SN, Southern Africa (Northern SN); SE, Southern Africa (Eastern); SW, Southern Africa (Western). The Tanzanian extant population haplotypes were mostly distributed into L3: haplotype 1, 3, 5 and 6 in EA; haplotype 4 in CE. However, the distinctive haplotype 2 was in the CV haplotype from L2.

Examining the relative frequency of extant East African haplotypes in a phylogenetic context (Figure 5) clearly indicates a substantial loss of genetic diversity compared to historic samples, but the remaining haplotypes span multiple lineages within in the CE and EA haplogroups. The phylogenetic tree also confirms observational records that the animals that had been translocated from South Africa were originally of East African origin; however, the introduced haplotypes (3,5, and 6) were all closely related and from a single EA cluster. None of the haplotypes at high frequency in the southern region of Africa (red circles, Figure 5) were detected in the east Africa region. The tree also indicates that some of the diversity that has been lost in Tanzania has been found in modern samples from Kenya.

## Discussion

This study presents the first assessment of genetic diversity of extant black rhinoceros populations in Tanzania and a neighboring population in Maasai Mara, Kenya. The study adds to the assessment of the global distribution of mtDNA diversity described by Moodley et al. (2017) and provides critical information about genetic relatedness that can inform conservation management of the current rhinos and other wildlife populations within the region. Although the comprehensive study by Moodley et al. (2017) had included three “recent” samples from Tanzania, only one was obtained from an animal which could still be alive. Therefore, our study fills in a major gap in the knowledge about current genetic diversity of black rhinoceros in two East African countries. As predicted from the drastic bottleneck that the extant populations experienced, with founding by only a few individuals, we found that current nucleotide diversity within the extant mtDNA control region was substantially lower than extant samples from Kenya and historical samples from Tanzania. This suggests that the current population in Tanzania have lost genetic variation, due not only to the dramatic population decline across Africa (Moodley et al. 2017), but also the current strategies for population management. The lack of allele sharing between the native and translocated populations in the Serengeti ecosystem suggests that there has been limited dispersal, likely due to the intensive protection zone management that is currently in place, which does not allow animals to leave their current home range. This has likely reduced the potential impacts of previous translocations. Since we found that previous black rhino reintroductions have restored some of the lost diversity, rather than bringing in completely new genetic variation from unrelated populations, opening up corridors to dispersal could allow more mixing of genetic variants, without unknown risks of outbreeding depression. In addition, some of the haplotypes that had been lost from Tanzania are still present in extant Kenyan samples, opening up the possibility for cross-border management actions to provide a genetic “rescue” in both countries without introducing genetic variants from outside East Africa. Another intriguing finding is that we found a highly divergent haplotype that matches sequences identified as the *D. b. ladoensis* subspecies in Genbank; both our study and Moodley et al. (2017) detected this haplotype in the Maasai Mara (Kenya) and Moru (Institute), suggesting that subspecies status should be reconsidered in relation to management units, as it is not currently listed in the IUCN red list.

### Current levels of genetic variation in Tanzania

Despite the recent decline in black rhinoceros populations, a moderate haplotype diversity (0.7) has been maintained, which is consistent with findings in other regional populations such as those in Zimbabwe (0.76) and Kenya (0.88) (Muya et al. 2011; Moodley et al. 2017). Haplotype diversity, nucleotide diversity and the number of haplotypes are all influenced by the proportion of the population sampled (Goodall-Copestake et al. 2012); in our study we sampled varying proportions of each population but we had the advantage of knowing how many maternal lineages were expected, due to detailed information on the founders. For example, we sampled 84% of the current population from Moru but only two haplotypes (Haplotype 1, Haplotype 2) were identified, which is consistent with founding from two females. Nevertheless, this population had the highest nucleotide diversity among the populations sampled (pi = 0.017), which was comparable to pre-bottleneck historic patterns (pi=0.021). However, this was due to the presence of highly divergence haplotype that is most closely related to *D. b ladoensis*; the presence of highly distinctive haplotypes can give a distorted view of the genetic health of populations if relying only on nucleotide or haplotype diversity, particularly if only a few maternal lineages are represented. In the Maasai Mara population, we found three haplotypes (haplotype 1, 2 and 4; n=12), similar to what was found by Moodley (2017) with n=15. However, a study conducted by Muya et al (2011; Figure 2) found eight additional haplotypes in the Masai Mara population (including haplotypes 3 and 6); although they didn’t provide a detailed sample list, these appear to have been collected in the past decade, suggesting that diversity could be higher than our sampling suggested. Moodley et al. (2017) sequenced mtDNA for additional samples from the Masai Mara that had been used in the nuclear analysis by Muya et al. (2011) and also found haplotype 2 (as well as 1 and 4). We sampled all five individuals found in the other native populations in this region (Nyamalumbwa) but identified only a single haplotype (haplotype 1); sharing of this haplotype with all of the other native populations, as well as samples from Kenya (Moodley et. 2017; Githui et al. 2014), suggests that this could be an ancestral allele, reflecting historical, rather than recent connectivity among these populations. Moodley et al. (2017) also sampled historic individuals from Zambia that had haplotype 1, despite being classified as a different subspecies (*D. b. nyasae*). Whether this reflects admixture between subspecies or misclassification would require further investigation. However, The uneven distribution of haplotypes across these populations means that allowing movements between one population to another through natural dispersal could supplement diversity within the less diverse population, such as has been suggested for Bison herds in the USA and Canada to restore gene flow to achieve long-term species viability (Davies et al. 2022). On the other hand, managing which animals are allowed to move could also be desirable. For example, the relatively high presence of haplotype 1 could suggest over-representation of particular maternal lineages in the native populations (leading to increased inbreeding). Avoiding translocating or allowing movements of individuals with this mtDNA haplotype to populations where it is already present could be worthwhile and preferentially allowing individuals with rarer haplotypes to move could be more effective for restoring lost genetic diversity. Despite retaining highly differentiated haplotypes, the concern for sustainability of the current populations is the low number of maternal lineages confirmed by the mtDNA variation. This means that allowing movements of native individuals might not be enough to maintain sufficient genetic diversity. For example, the Moru population was formed by three native black rhinoceroses, two females who survived the poaching catastrophe and one male which migrated from Ngorongoro. Finding of only two maternal haplotypes from 32 sampled individuals suggests that both maternal lines have been retained, which is good. However, the risk of maintaining isolated populations that started with only a few founders should not be ignored. Mkomazi and Ngorongoro had the highest number of haplotypes (three each), including haplotypes that are shared with Ndasiata despite a large geographical distance which exists in between. This is likely because the Mkomazi and Ndasiata populations were formed by individuals from captive breeding programmes and Ngorongoro had a mix of native and reintroduced individuals (Fyumagwa & Nyahongo 2010). Managing these population as a single metapopulation rather than in IPZ would allow movements of individuals from Ngorongoro through to Maasai Mara and could allow supplementation of the native populations with additional genetic variation from past translocations.

### Differentiation between subpopulations in Tanzania

The AMOVA analysis revealed that a high percentage of variation exists among individuals within populations, but the overall differentiation was moderate, which could reflect historical sharing of alleles and movement between populations. In Kenya, Muya et al. (2011) found the highest Fst (0.729) between Chyulu and the Masai Mara, neither of which included introduced individuals. However, the Chyulu population had been bottlenecked to only two individuals, consistent with the presence of only two mtDNA haplotypes. Since the two populations are within the same ecosystem, this suggests recent restriction of movement, similar to in our study. The lack of sharing of the introduced haplotypes in geographically close populations suggests that there is more restriction of gene flow than their home ranges would predict. Black rhinoceros are solitary and highly mobile; their estimated home ranges in the Serengeti are between 40 and 133 km^2^, regardless of sex (Frame 1980). However, the Intensive Protection Zones (IPZs) strategy, with no any movement of rhinoceros allowed between populations (Fyumagwa & Nyahongo 2010), means that these home range sizes are not realized. A vivid example is the Ndasiata-Nyamalubwa population, which didn’t share any haplotypes with Nyamalumbwa, despite being located very close in the same Serengeti-Mara ecosystem (Figure 2 and Figure 4) had the highest Fst value (0.97). Observational data suggests that the animals would move further if left more unconstrained., For example, in the Ngorongoro Crater and Moru, on several occasions, rhinos, especially bulls, have left their IPZ in search of new habitat or to escape from territorial fights; likewise, Moru individuals have escaped to Mwiba-Makoa areas. However, the management requires pushing them back into their respective IPZ. For example, in 2004 a young bull from NCA was sited near lake Eyasi, 100km from NCA, but was immobilized and returned back (Fyumagwa & Nyahongo 2010).

### Phylogenetic context of Tanzanian haplotypes

In Tanzania, translocation or assisted dispersal has been used as a tool for dispersal of rhinoceroses between populations for the purpose of increasing diversity and gene flow. In previous years (1997-2018), a total of 23 black rhinoceroses were translocated to Tanzania from areas outside East Africa but this was done without consideration of genetic variation (Fyumagwa & Nyahongo 2010). Only three haplotypes (haplotypes 3, 5 and 6) were found from 13 reintroduced rhinoceroses sampled in this study. The parental stock of these individuals were captured from Kibodo area, Tsavo National Park, Isiolo and Tana river in Kenya between 1960 and 1980 and taken outside East Africa (**Error! Reference source not found.**) in highly protected areas such as zoos and closed sanctuaries as a measure to rescue them from poaching in the wild. Phylogenetic reconstruction of the mtDNA demonstrated the maternal lineages introduced to Ndasiata, Mkomazi and Ngorongoro were of East African origin, despite individuals being translocated from European zoos or captive population in South Africa. This was confirmed by the presence of all three reintroduced haplotypes occurring across multiple populations in both recent and historic samples from Kenya (Muya et al. 2011; Githui et al. 2017; Moodley et al. 2017). However, the three introduced haplotypes were closely related and clustered together on the tree, suggesting that the previous translocations achieved limited augmentation of genetic diversity in the extant populations. However, the presence of a wider range of the “lost” Tanzanian haplotypes in modern Kenyan samples suggests that translocations within East Africa could be more beneficial than more expensive and risky long-distance translocations. This also could be important to avoid translocation catastophes, which can occur when animals that have been kept under benign captive conditions are released to wild environments (Chipman et al. 2008).

The phylogenetic tree also suggested that mitochondrial control region sequence variation is highly structured, allowing careful consideration of which lineages could be more beneficial to reintroduce. Tanzanian extant populations haplotypes were distributed into all three haplogroups (CV, CE and EA) found in East Africa but there were notable gaps in the presence of particular clades that were present historically. On the other hand, it is encouraging that haplotype 2, which was unique from the others, and clustered under the CV group is present in both Kenya and Tanzania. The historical range of the subspecies it represents was east of the Shari-Logone River to East Africa (Moodley et al. 2017). In the current study, individuals with this haplotype were restricted only to Moru and Masai Mara. Therefore, individuals possessing this haplotype might be unique and worthy of focus for conservation. However, since mtDNA only reflects the maternal lineage, it is possible that this haplotype represents a hybrid whose mother came from the now extinct subspecies and a *D. b. michaeli* father. The next step for specifically identifying individuals for translocation will be to obtain perspectives based on the nuclear genome, not only to confirm the status of individuals with rare maternal haplotypes but to identify conservation units based on both maternal and paternal contributions. We are currently taking a whole genome sequencing approach to address this question, using a subset of the individuals used in this study.

### Conservation implications

Our study has shown that the Tanzanian black rhinoceros populations has lost substantial variation in the mtDNA from the recent population decline but still maintains moderate genetic diversity within the populations. Recently translocated populations (Mkomazi and Ndasiata) have restored some of haplotypes that were previously present. This diversity could be spread by allowing movement of rhinos between the populations or strategic translocation of individuals between sites on a more local basis than has been done previously. Genetic diversity is an important factor for wildlife populations to persist as high genetic diversity within a population is important for the long-term survival of the species (Caughley 1994; Frankham 1995). Hence, managing genetic diversity should be one of the primary goals in various black rhinoceros conservation initiatives and when developing an active conservation strategy for the Tanzanian black rhinoceros population. The revealed variation in populations that had been historically connected can also provide a useful tool in translocation/reintroduction strategies from both wild and captive breeding programs. However, as data from the mtDNA can only give us a picture of maternal lineages, the addition of whole genome sequence analysis is necessary to refine black rhinoceros management programmes in Tanzania and East Africa.

## Statements & Declarations

### Funding and acknowledgement

This work was supported by the following funders Zurich Zoo, National Geographic early career grants [Grant number EC-51221R-18], and the Paul Tudor Jones Family Trust.

We are grateful to the following institutions for making their collections available to this study: Tanzania National Parks (TANAPA), Tanzania Wildlife Authority (TAWA), Ngorongoro Conservation Authority (NCA) and the Kenyan Wildlife Service (KWS).

We thank Nelson Mandela African institute of science and Technology, Arusha, Tanzania and Kenya Wildlife Services for providing us a platform in their laboratories to perform DNA extraction of the samples.

Also, we thank Tanzania Wildlife Research Institute (TAWIRI) and Tanzania Commission for Science and Technology (COSTECH) for granting us permits to conduct the study.

### Author Contributions

Conceptualization: [Ronald Mellya]; Funding acquisition: [Barbara Mable, Grant Hopcraft, Ronald Mellya]; Data curation: [Ronald Mellya, Dickson Wambura, Emmanuel Macha, Moses Otiende, Iddrisa Chuma, Ernest Eblate]; Methodology: [Ronald Mellya, Barbara Mable]; Formal analysis: [Ronald Mellya, Barbara Mable]; Investigation: [Ronald Mellya, Ernest M. Eblate, Dickson Wambura, Emmanuel Macha, Moses Otiende, Iddrisa Chuma, Ernest Eblate, Elizabeth Kilbride, Grant Hopcraft Barbara Mable]; Project Administration: [Ronald Mellya, Grant Hopcraft and Barbara Mable]; Writing - original draft preparation: [Ronald Mellya]; Writing - review and editing: [Ronald Mellya, Ernest M. Eblate, Dickson Wambura, Emmanuel Macha, Idrissa Chuma, Grant Hopcraft, Barbara Mable]; Funding acquisition: [Grant Hopcraft], [Ronald Mellya]; Resources: [Barbara Mable, Grant Hopcraft, Moses Otiende, Ernest Eblate]; Supervision: [Barbara Mable].

### Corresponding authors

Correspondence to Ronald Mellya.

### Data Availability

The datasets generated during the current study are available on GenBank repository, accession numbers: OQ095383-OQ095388.

